# Wnt/beta-catenin signaling regulates Vascular Endothelial Growth Factor (VEGF) receptors in central nervous system endothelial cells

**DOI:** 10.64898/2026.09.03.745507

**Authors:** Yanshu Wang, Ningyu Zhu, Amir Rattner, Philip M. Smallwood, Jeremy Nathans

## Abstract

In CNS endothelial cells (ECs), VEGF signaling promotes vascular permeability, and Wnt/beta-catenin signaling reduces vascular permeability by controlling the gene expression program for the blood-brain barrier. Here we show, using genetic mosaics, that, in mouse brain ECs *in vivo*, an increase in Wnt/beta-catenin signaling produces an increase in VEGFR1 levels and a decrease in VEGFR2 levels, and a decrease in Wnt/beta-catenin signaling produces a decrease in VEGFR1 levels and an increase in VEGFR2 levels. As VEGFR1 functions as a decoy receptor to reduce VEGF signaling through VEGFR2, these data imply that Wnt/beta-catenin signaling acts at the receptor level to reduce VEGF signaling. In HEK/293T cells, VEGF signaling is suppressed by Wnt/beta-catenin signaling in a dose-dependent manner, with nearly complete suppression observed at levels of Wnt/beta-catenin signaling that produce little or no change in VEGF receptor levels. These data reveal two mechanisms by which VEGF signaling is regulated by Wnt/beta-catenin signaling.

## Introduction

Vascular Endothelial Growth Factor (VEGF) signaling via transmembrane VEGF receptors (VEGFRs) regulates endothelial proliferation, migration, permeability, and homeostasis (Pérez-Gutiérrez and Ferrara, 2023). In the context of developmental and physiologic tissue hypoxia, induction of *Vegf* gene expression by hypoxia-inducible factors (HIFs) in target tissues orchestrates a pro-angiogenic response that results in the appropriate delivery of oxygen and nutrients (Forsythe et al., 1996; Semenza, 2003). An analogous VEGF-driven angiogenic response is observed in the context of pathologic neovascularization, as seen, for example, in neovascular age-related macular degeneration (AMD), proliferative diabetic retinopathy, corneal neovascularization, rheumatoid arthritis, and within solid tumors (Apte et al., 2019; Pérez-Gutiérrez and Ferrara, 2023).

VEGF was first identified and purified based on its ability to increase vascular permeability, and, as a consequence, it was originally named Vascular Permeability Factor (VPF) (Senger et al., 1983; Nagy et al., 2008). In central nervous system (CNS) vasculature, vascular permeability is restricted by the blood-brain barrier (BBB) and the blood-retina barrier (BRB). In neovascular AMD and diabetic retinopathy, increased vascular permeability, leading to retinal edema, is commonly observed and is attributed primarily to the effect of elevated tissue-derived VEGF acting on VEGFR2 in vascular endothelial cells (ECs) (Campochiaro, 2015). A current mainstay of therapy for neovascular AMD and diabetic retinopathy is intra-ocular injection of antibodies or antibody derivatives that bind to VEGF with high affinity (Campochiaro et al., 2016; Pérez-Gutiérrez and Ferrara, 2023).

In the CNS, angiogenesis and vascular barrier development and maintenance are also under the control of other signaling systems, including beta-catenin signaling (also referred to as canonical Wnt/ signaling; Rattner et al., 2022; Gastfriend and Daneman, 2026). In this system, ligands Wnt7a, Wnt7b, and Norrin are produced predominantly by glia and they act on Frizzled receptors, Lrp5 and Lrp6 coreceptors, and the ligand-specific co-activators Gp124 and Reck (for Wnt7a and Wnt7b) and Tspan12 (for Norrin) that are present on the plasma membrane of ECs (Xu et al., 2004; Stenman et al. 2008; Liebner et al., 2008; Daneman et al., 2009; Junge et al., 2009; Zhou et al., 2014; Vanhollebeke et al., 2015; Cho et al, 2017; Wang et al., 2018; Wang et al., 2020). In CNS ECs, beta-catenin signaling is required throughout life to maintain the BBB/BRB program of gene expression (Wang et al., 2012; Zhou et al., 2014). The phenotypic manifestations of this program in CNS ECs include (a) elimination of intercellular diffusional pathways, (b) down-regulation of transcytosis, (c) production of plasma membrane transporters for essential small molecules such as amino acids and glucose, and (d) production of ATP-driven pumps that extrude xenobiotics (Daneman and Prat, 2015).

Within the CNS, the vasculature that supplies the circumventricular organs (CVOs), represents an exception to the BBB program of differentiation (Morita and Miyata, 2012; Morita et al., 2016). The CVOs are small midline structures in which neurons have diffusional access to serum, either for the purpose of monitoring serum constituents (sensory CVOs) or for secretion of bioactive molecules into serum (secretory CVOs). In keeping with these functions and in contrast to BBB+ CNS capillaries, CVO capillaries exhibit high permeability, with a high density of fenestrations, high levels of the fenestration structural protein plasmalemma vesicle-associated protein (PLVAP), and an absence of classic blood-brain barrier (BBB) markers such as the glucose transporter GLUT1. CVOs also exhibit a high vascular density and ongoing vascular remodeling, characteristics that depend on VEGF signaling (Furube et al., 2014; Morita et al., 2015).

In earlier work, we and others showed that CVO ECs normally exhibit little or no beta-catenin signaling, and that genetically activating beta-catenin signaling in ECs converts high permeability BBB-CVO ECs to a low permeability BBB+ state (Benz et al. 2019; Wang et al., 2019). Furabe et al. (2014) and Morita et al. (2015) showed that CVOs are characterized by high levels of VEGF signaling, with neurons and glia producing VEGF and ECs expressing VEGFR2. Furabe et al. (2014) further showed that blocking VEGF signaling with a VEGFR2 antagonist reduces EC proliferation and vascular density in one CVO, the posterior pituitary. Pharmacologic blockade of VEGFR2 signaling also reduces the physiologic action of peripherally applied leptin, which likely reflects reduced vascular permeability in a second CVO, the median eminence (Jiang et al., 2020). In the retina and in EC monolayers *ex vivo*, beta-catenin signaling reduces barrier leakage in the context of a VEGF-induced increase in permeability (Díaz-Coránguez et al., 2020). Taken together, these data are consistent with a model in which beta-catenin and VEGF signaling interact in a mutually antagonistic fashion in ECs to regulate vascular permeability.

In the present study, we have explored regulatory interactions between beta-catenin signaling and VEGF signaling. Cross-regulatory interactions have been observed among several pathways that modulate ECs, including the Notch, VEGF, and Tie2/Angiopoietin pathways (Adams and Alitalo 2007; Jakobsson et al., 2009; Geudens and Gerhardt, 2011; Potente et al., 2011). Beta-catenin signaling has been implicated as a positive regulator of VEGF-A and VEGF-C expression (Zhang et al., 2001; Skurk et al., 2005; Kazanskaya et al., 2008; Gore et al., 2011), but the regulation of VEGFRs by beta-catenin signaling has remained largely unexplored. Here we show, in CNS ECs *in vivo*, that beta-catenin signaling leads to increased levels of VEGFR1, a decoy receptor, and reduced levels of VEGFR2. We further show, in HEK/293T cells, that beta-catenin signaling supresses signaling by VEGFR2. These data suggest that the presence of beta-catenin signaling in BBB+ ECs contributes to the low permeability of BBB+ ECs by inhibiting VEGF signaling and, conversely, that the low level of beta-catenin signaling in CVO ECs contributes to the high permeability state of the CVO vasculature by licensing VEGF signaling.

## Results

### Beta-catenin signaling regulates VEGFR1 and VEGFR2 levels in brain vasculature

As a first step in exploring the question of whether beta-catenin signaling regulates VEGFR levels in ECs, we asked whether BBB-CVO ECs (with low levels of beta-catenin signaling) exhibit levels of VEGFR1 or VEGFR2 that differ from the levels in surrounding BBB+ non-CVO ECs (with high levels of beta-catenin signaling). Immunostaining of the subfornical organ and the area postrema, two sensory CVOs, revealed a greater intensity of VEGFR2 immunostaining compared to the surrounding BBB+ vasculature (Figure 1A and B). In the area postrema, the immunostaining intensity of VEGFR1 was lower than the levels in the surrounding BBB+ vasculature (Figure 1B). Quantification of the relative VEGFR1 and VEGFR2 immunostaining intensities in ECs in and around the area postrema confirmed the impression obtained by visual inspection (Figure 1C).

**Figure 1.**
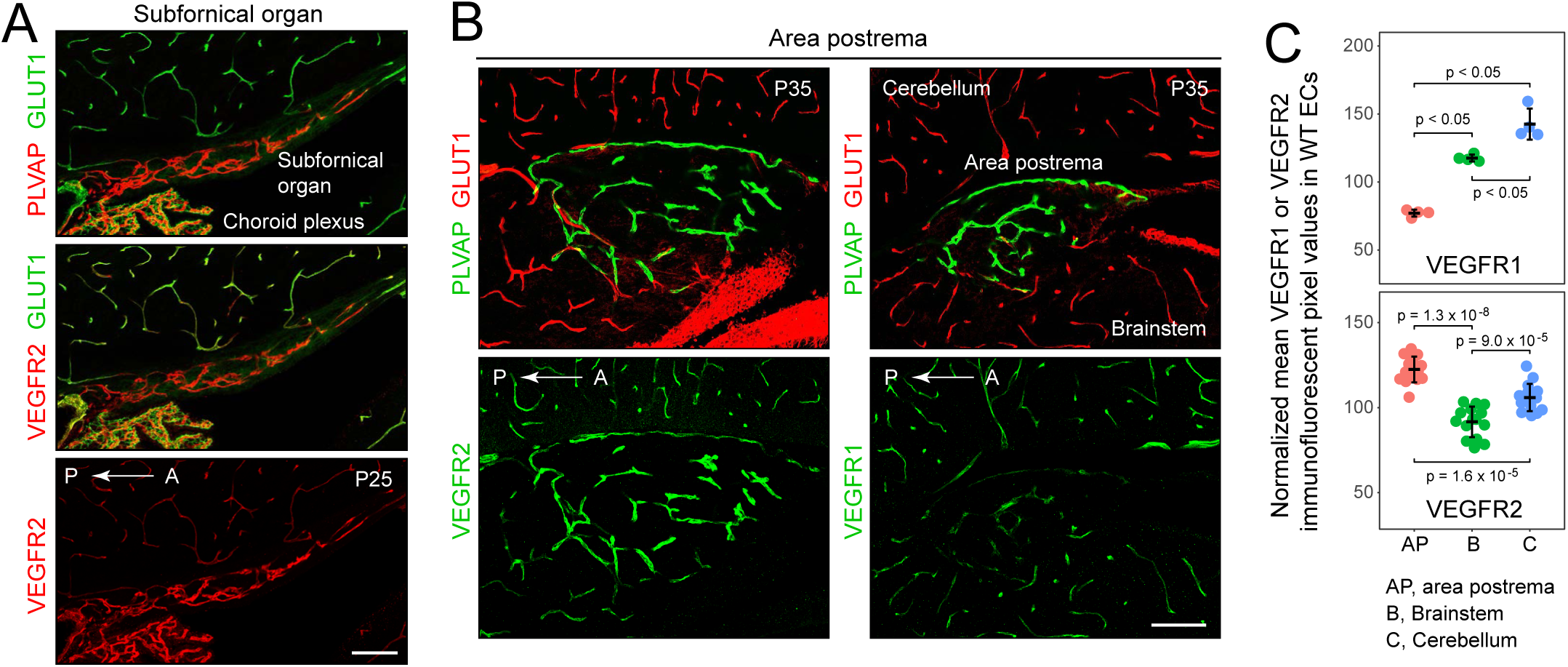
In WT mice, VEGFR1 levels are reduced and VEGFR2 levels are elevated in ECs in the area postrema relative to surrounding BBB+ ECs. (A) Sagittal section showing elevated VEGFR2 immunostaining in subfornical organ (SFO) ECs, which are GLUT1-/PLVAP+. The choroid plexus is in the upper right. A, anterior; P, posterior. The ages of the mice in postnatal days (P) are indicated in each set of panels in this and all other figures. Scale bar, 100 um. (B) Sagittal section showing elevated VEGFR2 and reduced VEGFR1 immunostaining in area postrema (AP) ECs, which are GLUT1-/PLVAP+. A, anterior; P, posterior. Scale bar, 100 um. (C) Quantification of VEGFR1 and VEGFR2 immunofluorescent intensities in ECs in the AP, brainstem, and cerebellum in P35 mice. Each data point in this panel and in the panels in Figures 2-4 represents the mean value for the ECs in a single confocal image.

To further assess the effects of the BBB+ vs .BBB-state on VEGFR2 levels, we compared BBB+ vs. BBB-ECs in *Fzd4^-/-^* cerebella, where previous work had demonstrated that reduced EC beta-catenin signaling leads a subset of cerebellar ECs to adopt a BBB-fate, as determined by their conversion from GLUT1+/PLVAP-to GLUT1-/PLVAP+ (Wang et al, 2018). In comparisons between BBB- and BBB+ ECs in the same tissue sections of *Fzd4^-/-^* cerebella, we observed an increase in VEGFR2 immunostaining intensity in BBB-ECs compared to BBB+ ECs (Figure 2A and B).

**Figure 2.**
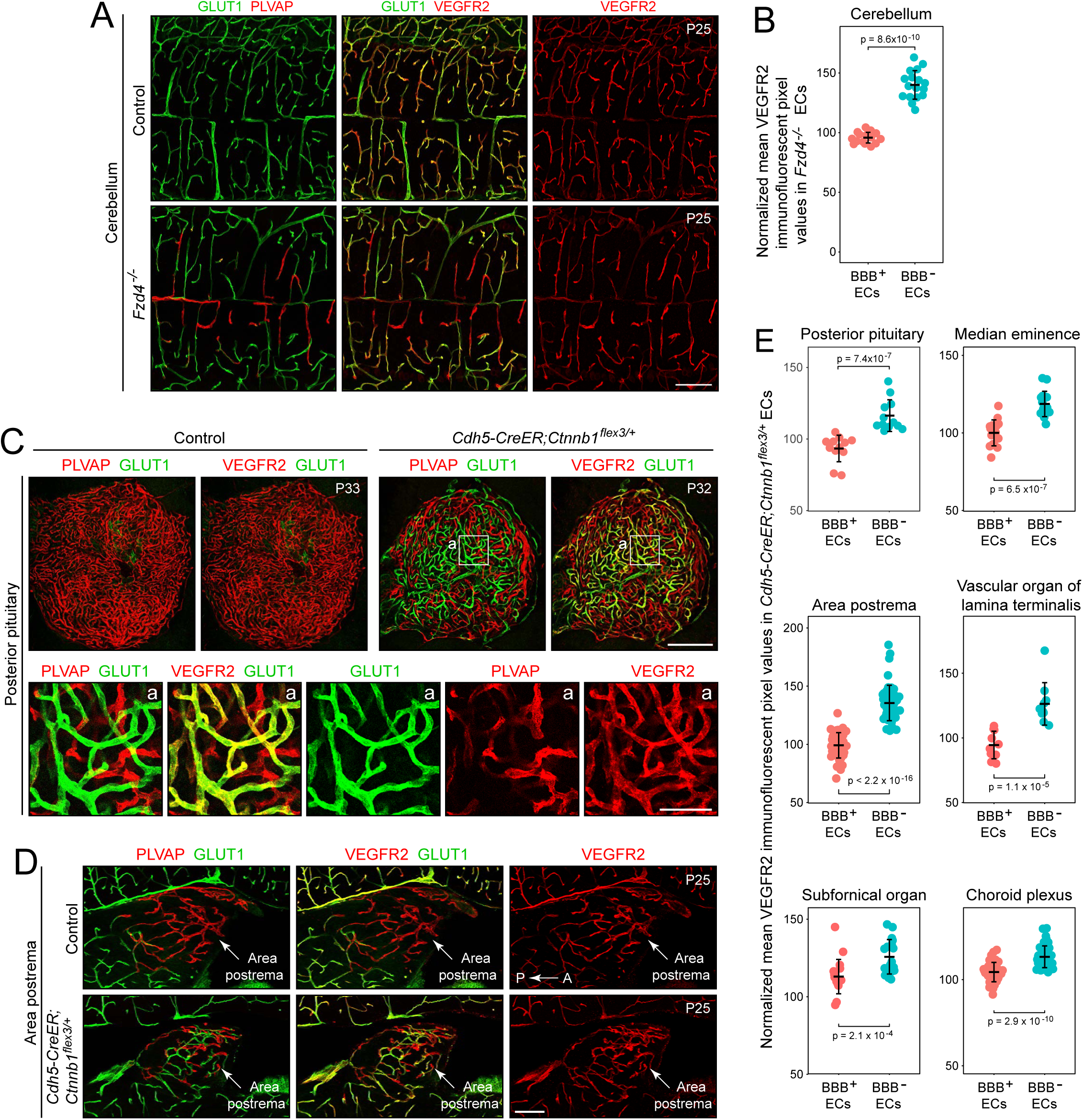
VEGFR2 levels are elevated in CNS ECs with reduced beta-catenin signaling and reduced in ECs with elevated beta-catenin signaling. (A) VEGFR2 immunostaining of the *Fzd4^-/-^* cerebellum, which exhibits a mixture of GLUT1-/PLVAP+ and GLUT1+/PLVAP-ECs. Scale bar, 100 um. (B) Quantification of VEGFR2 immunofluorescent intensities in ECs in images like the one in (A), with data points divided into BBB+ or BBB-ECs. (C,D) VEGFR2 immunostaining of *Cdh5CreER;Ctnnb1 ^flex3/+^* genetic mosaic vasculature in the posterior pituitary (C) and the area postrema (D), which exhibit a mixture of GLUT1-/PLVAP+ and GLUT1+/PLVAP-ECs. In (C), the region demarcated by white squares in the two upper right panels is enlarged in the lower set of five panels. Scale bars in (C), 200 um (upper panels) and 50 um (lower panels). Scale bar in (D), 100 um. (E) Quantification of VEGFR2 immunofluorescent intensities in ECs in *Cdh5CreER;Ctnnb1 ^flex3/+^* genetic mosaic vasculature in five CVOs and in the choroid plexus at P25-P35, with data points divided into BBB+ or BBB-ECs, as in (B).

The comparison described above between CVO and non-CVO ECs is suggestive of a role for beta-catenin signaling in modulating VEGFR levels, but it does not rule out effects from other developmental or homeostatic pathways. We therefore used a tissue mosaic paradigm to assess beta-catenin gain-of-function effects on VEGFR levels. Specifically, we genetically stabilized beta-catenin, the protein product of the *Ctnnb1* gene, in a subset of *Cdh5CreER;Ctnnb1 ^flex3/+^*ECs by treating mice with a low dose of 4-hydroxytamoxifen (4HT). [In the *Ctnnb1 ^flex3^* allele, the third exon of the *Ctnnb1* gene is flanked by *loxP* sites. Cre-mediated excision of this exon removes a beta-catenin phosphorylation site that is required for ubiquitinylation and proteosomal degradation, creating a small in-frame deletion that leaves the beta-catenin protein otherwise unperturbed (Harada et al., 1999).] In the genetically mosaic vasculature of the CVOs, VEGFR2 immunostaining intensity was quantified in GLUT1+/PLVAP-ECs and in GLUT1-/PLVAP+ ECs within the same tissue sections (Figure 2C-E). In each of the five CVOs examined – the posterior pituitary, median eminence, area postrema, vascular organ of the stria terminalis, and subfornical organ – VEGFR2 levels were lower in the GLUT1+/PLVAP-(i.e., BBB+) ECs than in GLUT1-/PLVAP+ (i.e., BBB-) ECs. The same pattern was also seen in genetically mosaic vasculature in the choroid plexus, the high permeability vasculature the generates the cerebrospinal fluid (Figure 2E).

*Cdh5CreER;Ctnnb1 ^flex3/+^*vascular mosaics were similarly analyzed for changes in VEGFR1 levels in GLUT1+/PLVAP-ECs and in GLUT1-/PLVAP+ ECs (Figure 3). In contrast to the results of the VEGFR2 analyses, in each of the five CVOs examined and in the choroid plexus, VEGFR1 levels were higher in the GLUT1+/PLVAP-ECs (i.e., BBB+ ECs) than in GLUT1-/PLVAP+ ECs (i.e., BBB-ECs) (Figure 3A and B). A smaller reduction in VEGFR1 levels was observed in GLUT1-/PLVAP+ (i.e., BBB-) ECs compared to GLUT1+/PLVAP-(i.e., BBB+) ECs in the *Fzd4^-/-^* cerebellum (Figure 3C).

**Figure 3.**
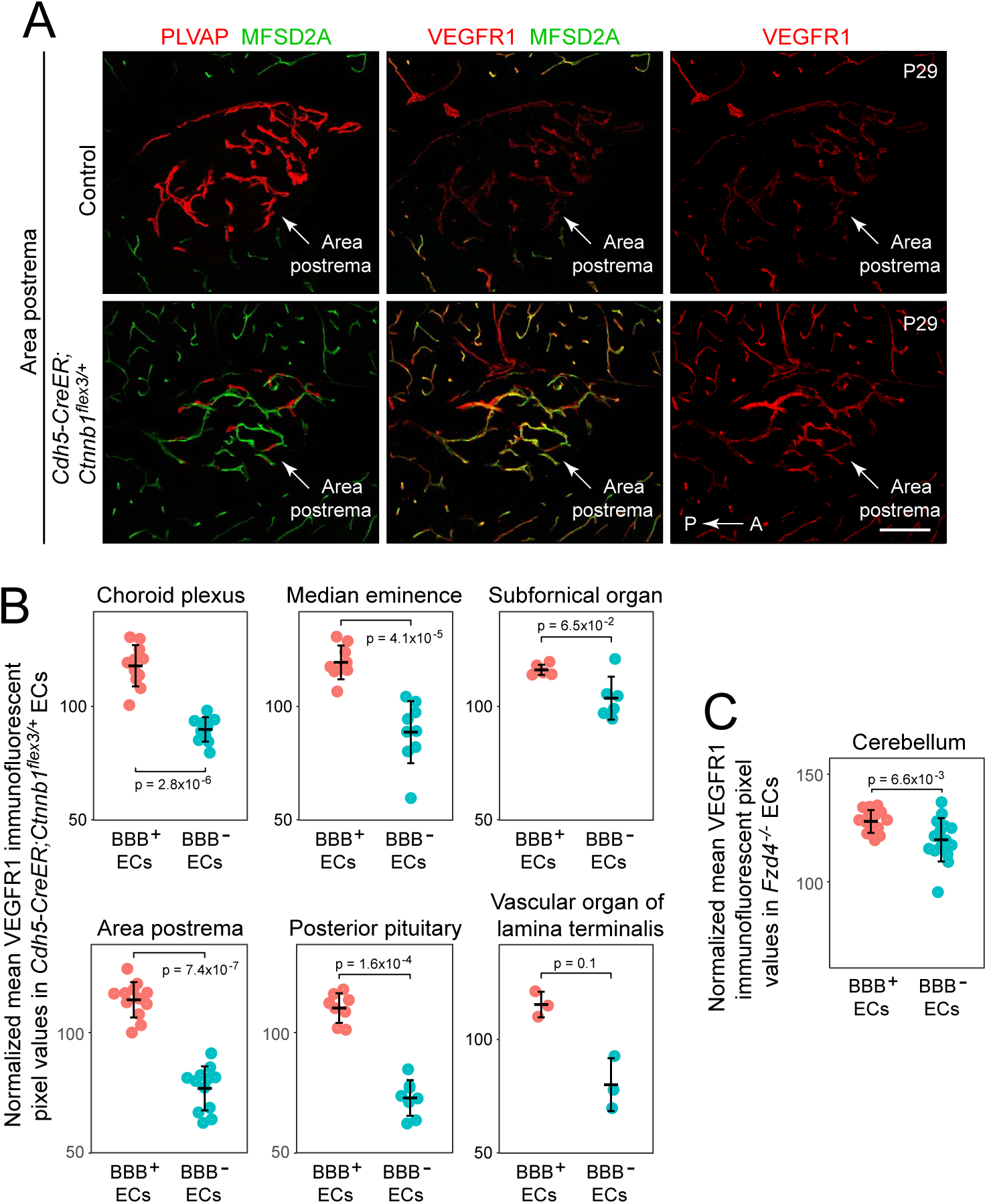
VEGFR1 levels are reduced in CNS ECs with reduced beta-catenin signaling and elevated in ECs with elevated beta-catenin signaling. (A) VEGFR1 immunostaining of *Cdh5CreER;Ctnnb1 ^flex3/+^* genetic mosaic vasculature in the area postrema, which exhibits a mixture of GLUT1-/PLVAP+ and GLUT1+/PLVAP-ECs. ECs in the WT area postrema are GLUT1-/PLVAP+. A, anterior; P, posterior. Scale bar, 100 um. (B) Quantification of VEGFR1 immunofluorescent intensities in ECs in *Cdh5CreER;Ctnnb1 ^flex3/+^* genetic mosaic vasculature in five CVOs and in the choroid plexus at P25-P35, with data points divided into BBB+ or BBB-ECs, as in Figure 2B. (C) Quantification of VEGFR1 immunofluorescent intensities in ECs in the *Fzd4^-/-^* cerebellum, with data points divided into BBB+ or BBB-ECs, as in Figure 2B.

By quantifying VEGFR2 levels in ECs in mice that are genetic mosaics (or that are genetically homogeneous but phenotypically mosaic, as in Figure 2A and B), these analyses controls for biological variables that could confound comparisons between mutant and wild type (WT) animals, such as differences in vascular anatomy, levels of tissue oxygenation, and signals from non-ECs. They also control for technical factors, such as differential tissue fixation, that might confound a comparison between mice.

Taken together, these data show that in mouse brain ECs, an *increase* in beta-catenin signaling produces an *increase* in VEGFR1 levels and a *decrease* in VEGFR2 levels, and a *decrease* in beta-catenin signaling produces a *decrease* in VEGFR1 levels and an *increase* in VEGFR2 levels. As VEGFR1 function as a decoy receptor to reduce signaling through VEGFR2, these data imply that beta-catenin signaling acts at the receptor level to reduce VEGF signaling.

### Effect of beta-catenin signaling on VEGFR1 and VEGFR2 levels in retinal vasculature

We next extended the genetic mosaic vascular analysis by asking whether VEGFR1 and/or VEGFR2 were regulated by beta-catenin signaling in the retinal vasculature, using low dose 4HT treatment of *Cdh5CreER*;*Fzd4^CKO/-^*mice to generate a mosaic of *Fzd4* WT and *Fzd4* KO ECs (Figure 4). In retina cross-sections and flat mounts, immunostaining for PLVAP (expressed by *Fzd4* KO BRB-ECs) and MFSD2A (expressed by *Fzd4* WT BRB+ ECs) revealed the EC mosaic. Immunostaining for VEGFR2 revealed expression in ECs, as well as in Muller glia, as previously described (Okabe et al., 2014).

**Figure 4.**
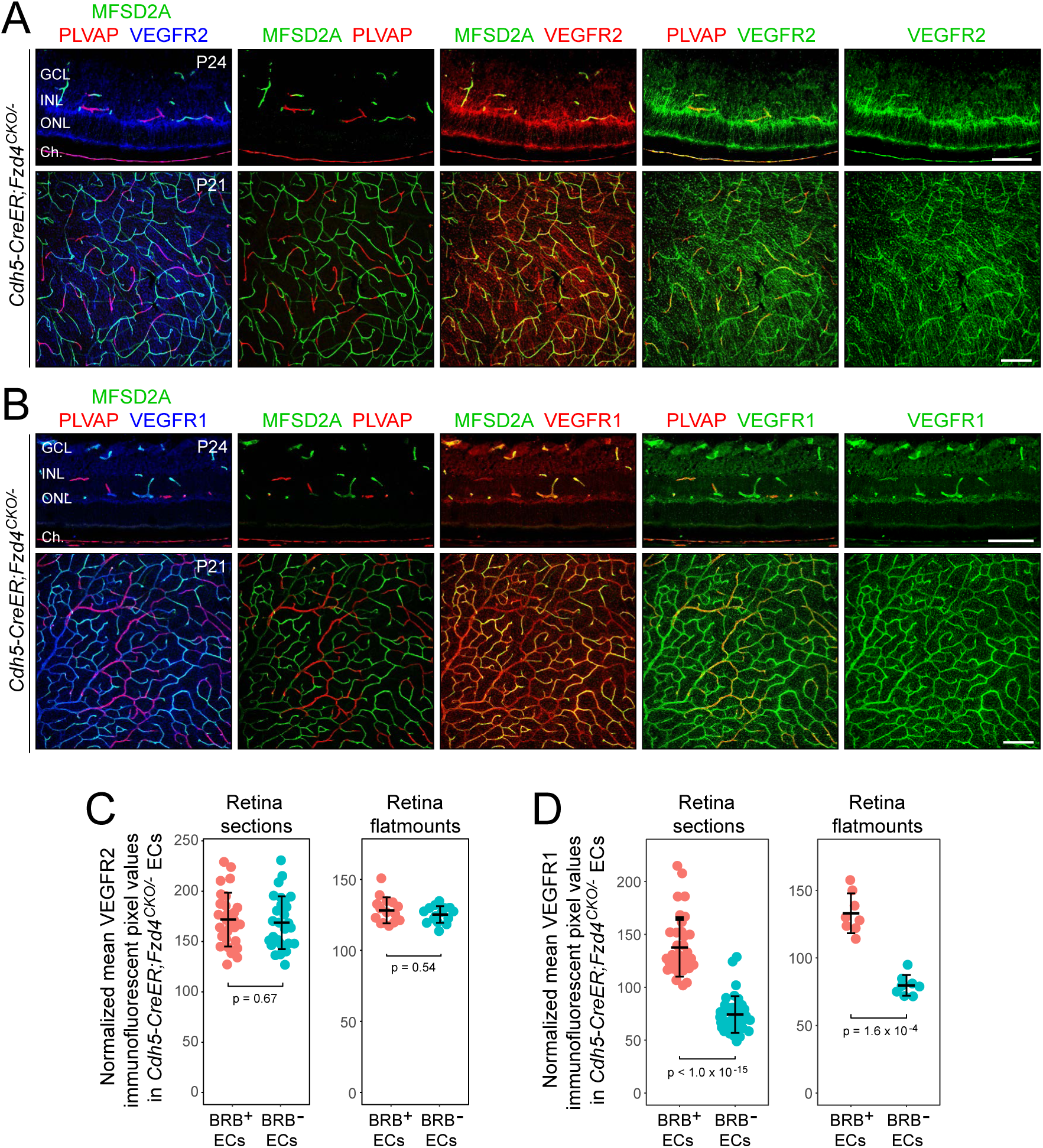
VEGFR1 and VEGFR2 levels in retinas with mosaic loss of beta-catenin signaling. (A,B) Cross-sections (upper) and flat mounts (lower) of retinas with *Cdh5CreER;Fzd4^CKO/-^* genetic mosaic vasculature, which exhibits a mixture of MFSD2A+/PLVAP-(i.e., BRB+) and MFSD2A-/PLVAP+ (i.e., BRB-) ECs. Tissues were immunostained for MFSD2A, PLVAP, and either VEGFR2 (A) or VEGFR1 (B). GCL, ganglion cell layer; INL, inner nuclear layer; ONL, outer nuclear layer; Ch, choroid. Retina cross-section scale bar for (A), 50 um. Retina cross-section scale bar for (B), 100 um. Retina flat mount scale bars for (A) and (B), 100 um. (C,D) Quantification of VEGFR2 (C) and VEGFR1 (D) immunofluorescent intensities in ECs in *Cdh5CreER;Fzd4^CKO/-^* genetic mosaic vasculature determined from retina cross-sections at P24 and from retina flat mounts at P21, with data points divided into BRB+ or BRB-ECs, as in Figure 2B.

Muller glial expression of VEGFR2 had the unfortunate effect of partially obscuring the VEGFR2 signal in ECs (Figure 4A). Quantification of VEGFR2 levels in BRB+ (i.e., *Fzd4* WT) and BRB-(i.e., *Fzd4* KO) ECs, showed no apparent difference (Figure 4C), although the high background of Muller glia-derived VEGFR2 makes this conclusion less certain compared to the analyses in the brain, where VEGFR2 is largely confined to ECs (Figures 1-3). Immunostaining for VEGFR1 revealed expression in ECs and a low background in non-ECs (Figure 4B). Quantification of VEGFR1 levels showed a ∼2-fold reduction of VEGFR1 in BRB-ECs compared to BRB+ ECs (Figure 4D), consistent with the observations in brain vasculature described above.

As an adjunct to the retinal EC mosaic analysis, the distribution and abundance of VEGFR2 was compared between (1) WT retinas, (2) two types of mutant retinas that have defective beta-catenin signaling in ECs [*Ndp^KO^* (*Ndp* codes for Norrin, the Frizzled4 ligand) and *Fzd4^KO^*] and (3) retinas with EC-specific knockout of *Transforming Growth Factor-Beta Receptor-1* (*Cdh5CreER*; *Tgfbr1^CKO/-^*).

Retinas with EC-specific KO of TGF-beta signaling exhibit an impoverished retinal capillary plexus and multiple tufts/clusters of endothelial cells near the vitreal face of the retina, similar to those seen in *Ndp^KO^* and *Fzd4^KO^*retinas (Xu et al., 2004; Ye et al., 2009; Zarkada et al., 2021; Wang et al., 2025; Figure S1). The overall anatomy of the retinal vasculature, visualized with PECAM1 immunostaining, and of the radial arteries, visualized with smooth muscle actin (SMA) immunostaining, was compared for the three classes of retinas (Figure S1). Compared to WT and EC-specific *Tgfbr1* KO retinas, *Ndp^KO^* retinas show modestly elevated VEGFR2 immunostaining in the hypertrophied surface vasculature and lower VEGFR2 immunostaining outside of the vasculature, principally in Muller glia (Figure S1A). Cross-sections of *Ndp^KO^*, *Fzd4^KO^*, and *Cdh5CreER*; *Tgfbr1^CKO/-^* retinas confirm the elevated level of VEGFR2 immunostaining in the EC tufts/clusters, as well as modestly reduced VEGFR2 levels in Muller glia compared to WT controls. However, the VEGFR2 levels in these retinas are difficult to quantify and interpret because the anatomy of the mutant retinal vasculature differs so greatly from that of the WT retina. Therefore, our principal conclusions from the comparisons in Figure S1 is that the optimal experimental paradigm for assessing the role of beta-catenin signaling on VEGFR levels in CNS ECs *in vivo* is a comparison between BBB+/BRB+ ECs and BBB-/BRB-ECs in a mosaic vasculature, in which case the vascular anatomy matches that of the WT control and the comparisons are being made between genetically distinct ECs in the same tissue section.

### Beta-catenin signaling and VEGFR2 signaling in HEK/293T cells

To investigate the interplay between beta-catenin signaling and VEGF signaling in a system that permits direct measurements of signal strength and experimental control over receptor and ligand levels, we turned to transient transfection of HEK/293T cells and an HEK/293 derivative stably transfected with the Super-Top-Flash (STF) reporter of beta-catenin signaling (Figure 5; Xu et al., 2004). In the absence of added receptors and/or ligands, these cells exhibit extremely low levels of VEGF and beta-catenin signaling. Transfection with a Wnt1 expression plasmid at 0.08-1.25 ng/well in a 96-well plate produced a dose-dependent increase in beta-catenin signaling, as measured by STF luciferase activity (Figure 5A). Transfection of a VEGFR2 expression plasmid and addition of VEGF to the growth medium – which together produced a dose-dependent increase in VEGF signaling, as measured with a NFAT-firefly luciferase reporter (Figure 5B, left panel) – had no effect on beta-catenin signaling, whether stimulated by Wnt1 plasmid co-transfection or by the addition to the medium of L6F4-2, an engineered antibody agonist that activates beta-catenin signaling by binding Frizzled4 and Lrp5, both of which are expressed at low levels in HEK/293T cells (Figure 5A and Figure 5 – figure supplement 1; Ding et al, 2023).

**Figure 5.**
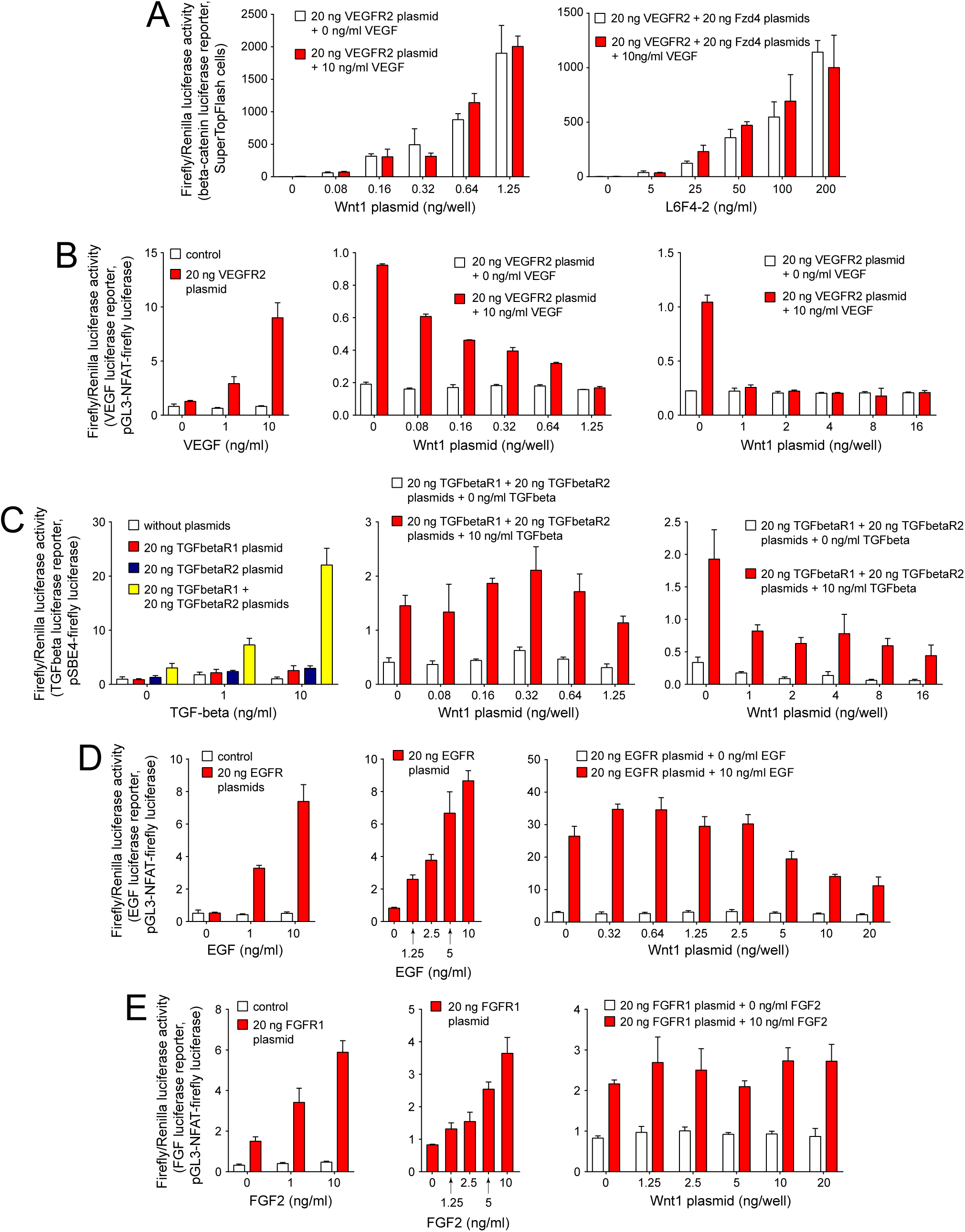
Effects of beta-catenin signaling on VEGF, TGF-beta, EGF, and FGF signaling in transfected HEK/293T cells, measured with luciferase reporters. (A) Lack of an effect of VEGF-VEGFR2 signaling on beta-catenin signaling stimulated by Wnt1 plasmid transfection (left). Lack of an effect of VEGF-VEGFR2 signaling on beta-catenin signaling stimulated by the Norrin mimic L6F4-2 (right). All experiments in this figure were carried out in triplicate in 96-well plates. (B) Calibrating VEGF-VEGFR2 signaling (left). Dose-dependent suppression of VEGF-VEGFR2 signaling with low levels of beta-catenin signaling (0.08-1.25 ng/well of Wnt1 plasmid) and complete suppression of VEGF-VEGFR2 signaling with high level beta-catenin signaling (1-16 ng/well of Wnt1 plasmid). (C) Calibrating TGFbeta-TGFBR1/TGFBR2 signaling and dependency on both receptor subunits (left). Little or no suppression of TGFbeta signaling with lower levels of beta-catenin signaling (0.08-1.25 ng/well of Wnt1 plasmid) (center), and modest suppression of TGFbeta signaling with high level beta-catenin signaling (1-16 ng/well of Wnt1 plasmid) (right). (D) Calibrating EGF-EGFR signaling (left and center). No suppression of EGF signaling except at the highest levels of beta-catenin signaling (>2.5 ng/well of Wnt1 plasmid) (right). (E) Calibrating FGF-FGFR1 signaling (left and center). No suppression of FGF signaling even at the highest levels of beta-catenin signaling (up to 20 ng/well of Wnt1 plasmid) (right).

In contrast to the insensitivity of beta-catenin signaling to VEGF signaling, VEGF signaling was suppressed by beta-catenin signaling in transfected HEK/293T cells in a dose-dependent manner (Figure 5B). This signal fell monotonically with co-transfection of the Wnt1 expression plasmid in the range 0.08 to 1.25 ng/well, and it was reduced to the background level at Wnt1 plasmid concentrations above 1 ng/well (Figure 5B, right two panels).

To assess the specificity of Wnt1-induced suppression of signaling by VEGFR2 in transfected HEK/293T cells, we compared it to the effect of Wnt1 on signaling by transforming growth factor-beta (TGF-beta), epidermal growth factor (EGF), and fibroblast growth factor (FGF) receptors (Figure 5C-E). Like VEGF signaling, each of these signaling systems uses single-pass transmembrane protein kinase receptors. In HEK/293T cells transfected with the relevant receptors – TGFBR1+TGBR2 for TGFbeta signaling, EGFR for EGF signaling, and FGFR1 for FGF signaling – and the relevant luciferase reporters, the cognate ligands produced the expected receptor- and ligand-dependent increases in luciferase activity (Figure 5C-E, left panels). For TGF-beta signaling, there was little or no effect of co-transfected Wnt1 plasmid concentrations up to 0.64 ng/well, and then a 2-3-fold decrease in signaling with Wnt1 plasmid concentrations between 1 and 16 ng/ml (Figure 5C, right two panels). For EGF signaling, there was little or no effect of co-transfected Wnt1 plasmid concentrations up to 2.5 ng/well, and then a ∼2-3-fold decrease in signaling with Wnt1 plasmid concentrations between 5 and 20 ng/ml (Figure 5D, right panel). For FGF signaling, there was little or no effect of co-transfected Wnt1 plasmid concentrations up to 20 ng/well (Figure 5E, right panel). Thus, among these four signaling systems, the order of sensitivity to Wnt1 inhibition is VEGF>TGFbeta>EGF>>FGF.

### High-level beta-catenin signaling broadly decreases receptor levels in transfected HEK/293T cells

The simplest mechanism by which beta-catenin signaling could suppress VEGF signaling in HEK/293T cells is by receptor down-regulation. To explore this possibility, we analyzed the levels of VEGFR2, TGFBR1, TGFBR2, EGFR, and FGFR1 (each expressed from transfected plasmids) by immunoblotting following co-transfection with different amounts of Wnt1 plasmid or without Wnt1 plasmid (Figure 5 – figure supplement 2). In these experiments, HEK/293T cells were grown in 6-well plates, which have 30-fold greater surface area/well than 96-well plates. Thus, normalized to surface area, 1 ng/well of Wnt1 plasmid in a 96-well plate, an amount that produced complete inhibition of VEGF signaling (Figure 5B), corresponds to 30 ng/well of Wnt1 plasmid in a 6-well plate. The immunoblots in Figure 5 – figure supplement 2 show that 30 ng/well of Wnt1 plasmid produced a ∼2-fold reduction in VEGR2 levels and had minimal effect on the levels of the other receptor types; 100 ng/well of Wnt1 plasmid produced a ∼2-fold reduction in the levels of all receptors except TGFBR2, which showed little change; and 300 ng/well of Wnt1 plasmid produced a several-fold reduction in the levels of all five receptors, with VEGFR2 showing the greatest effect. These experiments show that high levels of beta-catenin signaling lead to a general down-regulation of all five receptors. As this broad receptor down-regulation occurs at Wnt1 plasmid levels at least 10-fold higher than the levels that produce the dose-dependent suppression of VEGF signaling (Figure 5B), it is unlikely to account for the dose-dependent inhibition of VEGF signaling by beta-catenin signaling in HEK/293T cells, which likely reflects post-receptor regulation.

## Discussion

The experiments presented here reveal a previously undescribed link between beta-catenin signaling and VEGF signaling. In brain ECs beta-catenin signaling lowers the level of the signaling receptor VEGFR2 and raises the level of the decoy receptor VEGFR1, and in retinal ECs beta-catenin signaling raises the level of VEGFR1. These changes would be predicted to reduce VEGF signaling. Since VEGF signaling increases vascular permeability, a reduction in VEGF signaling should enhance vascular barrier function. In the CVOs, where vascular permeability is high, beta-catenin signaling in ECs is suppressed (Benz et al., 2019; Wang et al., 2019) and VEGF signaling in ECs is active (Furube et al., 2014; Morita et al., 2015). Our experiments show that in CVO ECs, suppression of beta-catenin signaling raises the level of VEGFR2 and lowers the level of VEGFR1, effects that would be predicted to enhance VEGF signaling.

At present, there is no system for quantifying VEGF signal strength at single cell resolution *in situ*, and, therefore, we have inferred the physiological consequences of changes in VEGFR levels based on the known functions of VEGFR1 and VEGFR2.

In our experiments with HEK/293T cells, we observed that beta-catenin signaling produced a dose-dependent suppression of VEGF signaling, and this suppression was more sensitive to beta-catenin signaling than was suppression of TGF-beta, EGF, and FGF signaling. More specifically, the dose-dependent suppression of VEGF signaling occurred in response to levels of Wnt1 plasmid that generated a steep increase in beta-catenin signaling. Unexpectedly, the levels of VEGFR2 protein did not change within this range of Wnt1 plasmid concentrations but only showed a reduction at Wnt1 plasmid concentrations ∼10-fold higher. Our current interpretation of the HEK/293T experiments is that (1) the reduction in VEGF signaling mediated by low concentrations of Wnt1 plasmid likely represents physiological cross-regulation that does not involve VEGFR2 down-regulation but instead functions downstream of VEGFR2, and (2) the reduced level of VEGFR2 that was observed by immunoblotting in response to 10-100-fold higher levels of Wnt1 plasmid co-transfection likely represents a response distinct from the Wnt-1 mediated suppression of VEGF signaling revealed with the luciferase reporter.

Taken together, these data point to two levels of regulation of VEGF signaling by beta-catenin signaling: (1) control of VEGFR1 and VEGFR2 levels, as demonstrated in CNS ECs *in vivo*, and (2) control of VEGF signaling at some point between the receptor and the control of target gene transcription, as demonstrated in HEK/293T cells. It will be of great interest to determine whether the second level of regulation occurs in CNS ECs *in vivo*. In sum, these experiments reveal a new link between two signaling systems that together control angiogenesis and vascular permeability.

### Relevance for therapeutics that activate beta-catenin

The efficacy of anti-VEGF therapies in treating neovascular AMD and diabetic retinopathy has stimulated the search for modulators of other EC signaling pathways that could provide mechanistically distinct approaches to treating these diseases (Joussen et al., 2021; Nguyen et al., 2022b; Cao et al., 2023). In this context, beta-catenin signaling represents an attractive target for several reasons. Most obviously, beta-catenin activators would be expected to enhance the expression of the genetic program for BRB maintenance and thereby counter-act disease-associated reductions in vascular barrier integrity; second, extrapolating from the anti-inflammatory action of EC beta-catenin signaling in the context of experimental auto-immune encephalitis (Lengfeld et al., 2017), beta-catenin activators might reduce the inflammation associated with retina vascular diseases (Ambati et al., 2013; Antonetti et al., 2021); and third, the observation that Norrin and a synthetic Norrin mimic suppress VEGF-induced retinal vascular permeability in rat and rabbit models and that Norrin suppresses VEGF-induced permeability in EC monolayers *in vitro* (Díaz-Coránguez et al., 2020; Nguyen et al. 2022a) suggests that beta-catenin activators might suppress pathologic VEGF signaling in the context of human retinal disease. The present study lends support to the third mechanism. We note that these three possible mechanisms of disease modulation – strengthening the BRB, reducing vascular inflammation, and inhibiting VEGF signaling – are not mutually exclusive, and it is possible that they could operate simultaneously and in a synergistic manner.

For clinically relevant activation of beta-catenin signaling, the challenge – until recently – was the lack of drug-like molecules. However, the discovery that beta-catenin signaling depends on the induced clustering of receptors (Frizzled) and co-receptors (Lrp5/6) paved the way for the development of engineered inducers of this pathway (Cong et al., 2004; Holmen et al., 2005; Janda et al., 2017). In particular, immunoglobulin derivatives that bring Frizzled4 and Lrp5/6 together exhibit potent bioactivity *in vitro* and *in vivo* (Tao et al., 2019; Chen et al., 2020; Chidiac et al., 2021; Ding et al., 2023; Post et al., 2023; Zhang et al., 2023). One of these inducers is currently in clinical trials for the treatment of diabetic macular edema, neovascular AMD, and branch retinal vein occlusion (https://clinicaltrials.gov; NCT06571045; NCT06957080, NCT07205887).

In the present work, we observed, in mosaic retinas, that loss of EC beta-catenin signaling lowered VEGFR1 levels but had little effect on VEGFR2 levels. However, unlike the CVO vasculature, where an increase in beta-catenin signaling can be monitored by immunostaining for GLUT1 and PLVAP (Figure 2), in non-CVO (BBB+ or BRB+) vasculature there is currently no method for marking ECs that have acquired a further increase in beta-catenin signaling. Thus, our genetic mosaic analysis of EC beta-catenin signaling in the retina was limited to loss-of-function experiments. Given our observation of lower VEGFR2 in CVO ECs with stabilized beta-catenin, it is possible that during treatment with a beta-catenin signaling agonist the resulting supra-physiologic levels of signaling in retinal ECs could lower VEGFR2. Similarly, our observation of elevated VEGFR1 in CVO ECs with stabilized beta-catenin suggests that supra-physiologic levels of beta-catenin signaling in retinal ECs might elevate VEGFR1.

Finally, if the post-receptor physiologic suppression of VEGF signaling by beta-catenin signaling that we observed in HEK/293T cells also applies to CNS ECs, that might constitute an additional mechanism by which pharmacologic elevation of EC beta-catenin signaling could reduce angiogenesis and vascular permeability in the context of retinal disease.

## Methods

### Mice

The following mouse lines were used: *Cdh5CreER* (Monvoisin et al., 2006); *Fzd4^CKO^* (Ye et al., 2009; JAX 011078); *Tgfbr1^CKO^* (Larsson et al., 2001; JAX 028701); *Ndp^KO^*(Ye et al., 2009; JAX 012287); *Fzd4^KO^* (Wang et al., 2001; JAX 012823); and *Ctnnb1^flex3^* (Harada et al., 1999). *Ndp* is located on the X-chromosome and therefore we refer to both female *Ndp^-/-^* and male *Ndp^-/Y^* mice as *Ndp^KO^*. All mice were housed and handled according to the approved Institutional Animal Care and Use Committee protocol of the Johns Hopkins Medical Institutions (protocol MO25M305).

### 4-hydroxytamoxifen preparation and administration

4HT (Sigma-Aldrich H7904-25MG) was dissolved at 20 mg/ml in ethanol by extensive vortexing. Sunflower seed oil (Sigma-Aldrich S5007) was added to dilute the 4HT to 2 mg/ml, and aliquots were stored at −80°C. Thawed aliquots were mixed well before injections. All injections were performed intraperitoneally.

### Antibodies

The following antibodies were used for tissue immunofluorescence: rat mAb anti-mouse PLVAP/ MECA-32 (BD Biosciences 553849); Rabbit mAb anti-mouse Mfsd2a (clone E8U2O, Cell Signaling Technology 80302); rabbit mAb anti-human GLUT1 (clone SA0377, Thermo Fisher Scientific MA5-31960); goat pAb anti-mouse VEGFR1/Flt-1 (R&D systems AF471); and goat pAb anti-mouse VEGFR2/KDR/Flk-1 (R&D systems AF644). Alexa Fluor-labeled secondary antibodies were from Thermo Fisher Scientific.

The following antibodies were used for Western blots: rabbit mAb anti-VEGFR2 (D5B1, Cell Signaling Technology 9698); rabbit mAb anti-FGFR1 (D8E4, Cell Signaling Technology 9740); rabbit mAb anti-EGFR (D38B1, Cell Signaling Technology 4267); rabbit mAb anti-TGF-beta receptor I (F6L3I, Cell Signaling Technology 49728); rabbit mAb anti-TGF-beta receptor II (E5M6F, Cell Signaling Technology 41896); mouse mAb anti-beta actin (2D4H5, Proteintech 66009-1-Ig); goat anti-rabbit IgG (H+L), HRP conjugate (Thermo Fisher Scientific 31460); and goat anti-mouse IgG (H+L), HRP conjugate (Thermo Fisher Scientific 31430).

### Tissue processing and immunohistochemistry

Tissues were prepared and processed for immunohistochemical analysis as described (Wang et al., 2012; Zhou et al., 2014). In brief, mice were deeply anesthetized with ketamine and xylazine and then perfused via the cardiac route with 1% paraformaldehyde (PFA) in phosphate-buffered saline (PBS). Non-ocular tissues were dissected and dehydrated in 100% cold methanol overnight at 4°C. Tissues were re-hydrated the following day in 1× PBS at 4°C for at least 3 hours before embedding in 3% agarose. Tissue sections of 100–200 μm thickness were cut using a Leica vibratome.

For whole-mount retinas, intact eyes were immersion fixed in 1% PFA in PBS at room temperature for 1 hour before the retinas were dissected. For eye sections, enucleated eyes were imbedded in optimal cutting temperature compound (Tissue-Tek 4853) and frozen in dry ice. Embedded eyes were cut into 14 μm sections with a Zeiss cryostat and stored on glass slides at –80°C. For immunostaining, sections were warmed to room temperature, fixed in 1% PFA at room temperature for 30 min, and washed in PBS before pre-blocking.

Tissue sections or whole-mount retinas were permeabilized in PBSTC (1× PBS+1% Triton X-100+0.1 mM CaCl_2_) overnight at 4°C and then incubated overnight at 4°C with primary antibodies diluted in 1× PBSTC+7% normal goat or donkey serum. The sections were then washed at least six times with 1× PBSTC over the course of 6–8 hours and subsequently incubated overnight at 4°C with secondary antibodies diluted in 1× PBSTC+7% normal goat or donkey serum. The next day, the sections were washed at least six times with 1× PBSTC over the course of six hours and mounted in Fluoromount G (SouthernBiotech 0100-01).

### Confocal microscopy

Confocal images were captured with a Zeiss LSM700 confocal microscope (20x or 40x objective) using Zen Black 2012 software, and processed with Image J, Adobe Photoshop, and Adobe Illustrator software. For experiments with control and mutant tissues, tissue processing, confocal imaging, and image processing were performed identically across genotypes unless stated otherwise.

### Quantification of VEGFR1 and VEGFR2 immunofluorescence intensities

For quantification of VEGFR1 and VEGFR2 immunofluorescence intensities, z-stacked confocal images were obtained by combining eight z-sections separated by two-micron intervals, for a total thickness of 16 microns. Image quantification was performed with Fiji/ImageJ (<u>fiji.sc</u>). In brief, each TIFF image was processed by first splitting the color channels [red channel (PLVAP), green channel (MFSD2A or GLUT1), and blue channel (VEGFR1 or VEGFR2)]. Thresholds were set for each channel to include large objects (ECs) but not background (i.e., objects consisting of only a few pixels). For consistency, the default threshold “Otsu” was used (https://imagej.net/ij/plugins/otsu-thresholding.html). Masks created from the red and green channels were used to extract the blue signal overlapping with each of these two channels. Three measurements were acquired from each image: (1) the mean pixel value of the blue channel overlapping with the red mask, (2) the mean pixel value of the blue channel overlapping with the green mask, and (3), for normalization, the mean pixel value of the blue channel across the entire image.

### Statistical analysis

All statistical values are presented as mean ± SD. The Wilcoxon rank sum test was used to measure statistical significance. Statistical tests were carried out using the following web sites: https://www.socscistatistics.com/tests/signedranks/default2.aspx and https://www.omnicalculator.com/statistics/wilcoxon-rank-sum-test#how-do-i-calculate-wilcoxon-rank-sum-test.

### Cell culture

HEK/293T cells and Super TOP Flash (STF) luciferase reporter cells (Xu et al, 2004) were maintained in DMEM/F-12 (Thermo Fisher Scientific 12500) supplemented with 10% fetal bovine serum at 37°C in a humidified incubator with 5% CO₂.

### Plasmids and STF cells

Reporter plasmids used in this study were: pGL3-NFAT luciferase (Addgene plasmid #17870) and pSBE4-Luc (Addgene plasmid #16495). The Super TOP Flash (STF) luciferase reporter cell line was used for Wnt/beta-catenin signaling assays (Xu et al, 2004). Receptor expression plasmids were pcDNA3.1-VEGFR2-V5/HIS (Addgene #205617), pcDNA3-ALK5/TGFBR1 (Addgene #80876), pRK5-TGFBR2-FLAG (Addgene #31719), pcDNA3.1-EGFR-V5/HIS (Addgene #201102), and pcDNA3.1-FGFR1c-V5/HIS (Addgene #201106).

### Recombinant proteins and reagents

Recombinant human EGF (R&D Systems 236-EG), recombinant human FGF2 (R&D Systems 3718-FB), recombinant human VEGF165 (R&D Systems 293-VE), and recombinant human TGF-beta1 (R&D Systems 7754-BH) were used as indicated. The sequence of L6F4-2, the Norrin-mimic antibody derivative (Ding et al., 2023), was derived from US patent 20230138045A1, heavy chain sequence ID #28 and light chain sequence ID #29. Heavy and light chain coding regions were synthesized by Integrated DNA Technologies (IDT; Coralville, IA) and inserted into pRK5. Large scale production and protein A purification of L6F4-2 from serum-free conditioned medium was performed by Antibodies, Inc (Davis, CA). Purified L6F4-2 was used at the indicated concentrations.

### Transfection

For luciferase assays, cells were transfected in 96-well plates using FuGENE HD Transfection Reagent (Promega E2311). Each well received a total of 80.5 ng DNA and 0.25 μl FuGENE HD. The DNA mixture included 0.5 ng Renilla luciferase plasmid as an internal control and 20 ng of each additional plasmid, including any firefly luciferase reporter plasmids. Salmon sperm DNA was used to adjust the total DNA amount to 80.5 ng per well. All transfections and luciferase assays were performed in triplicate. For Western blot experiments, cells were transfected using polyethyleneimine (PEI) as described below.

### Dual luciferase assays

2×10^4^ HEK/293T or STF cells/well were seeded in 96-well plates. The following day, the medium was replaced with fresh DMEM/F-12 containing 10% fetal bovine serum. Three hours later, cells were transfected as described above. After transfection, cells were cultured overnight and then changed to fresh medium containing the indicated ligand. Cells were incubated with ligand for 48 hours prior to luciferase measurement.

For Wnt/beta-catenin titration assays, Super TOP Flash (STF) cells were transfected with a CMV enhancer/promoter Wnt1 expression plasmid at 0, 0.08, 0.16, 0.32, 0.64, or 1.25 ng per well, as indicated. For NFAT luciferase assays, HEK293T cells were transfected with the pGL3-NFAT luciferase reporter plasmid together with the indicated receptor expression plasmids. All receptor plasmids were transfected at 20 ng/well. To confirm activation in each reporter system, VEGFR2-NFAT assays were first performed with VEGF at 0, 1, or 10 ng/ml, FGFR1-NFAT assays with FGF2 at 0, 1.25, 2.5, 5, or 10 ng/ml, and EGFR-NFAT assays with EGF at 0, 1.25, 2.5, 5, or 10 ng/ml. To determine whether Wnt1 modulated receptor-dependent NFAT reporter activity, cells were co-transfected with different amounts of Wnt1 plasmid and then stimulated with a fixed ligand concentration. In VEGFR2-NFAT assays, Wnt1 was transfected at 0, 0.08, 0.16, 0.32, 0.64, or 1.25 ng per well or, in a separate series, at 0, 1, 2, 4, 8, or 16 ng per well, followed by stimulation with 10 ng/ml VEGF. In FGFR1-NFAT assays, Wnt1 was transfected at 0, 1.25, 2.5, 5, 10, or 20 ng per well, followed by stimulation with 10 ng/ml FGF2. In EGFR-NFAT assays, Wnt1 was transfected at 0, 1.25, 2.5, 5, 10, or 20 ng per well, followed by stimulation with 10 ng/ml EGF. For SBE4-Luc assays, HEK293T cells were transfected with SBE4-Luc together with TGFBR1 alone, TGFBR2 alone, or both TGFBR1 and TGFBR2 expression plasmids. To confirm reporter activation, cells were stimulated with TGF-beta1 at 0, 1, or 10 ng/ml. To determine whether Wnt1 affected TGF-beta-dependent SBE4-Luc activity, cells were co-transfected with Wnt1 plasmid at 0, 0.08, 0.16, 0.32, 0.64, or 1.25 ng per well or, in a separate series, at 0, 1, 2, 4, 8, or 16 ng per well, followed by stimulation with 10 ng/ml TGF-beta1. To the effects of VEGF on beta-catenin signaling, STF cells were transfected with VEGFR2 plasmid at 20, ng/well and Wnt1 plasmid at 0, 0.08, 0.16, 0.32, 0.64, or 1.25 ng per well and then treated with 0 or 10 ng/ml VEGF. As a second assessment of the effects of VEGF on beta-catenin signaling, STF cells were transfected with Fzd4 and Lrp6 expression plasmids (20 ng each) and treated with L6F4-2 at 0, 5, 25, 50, 100, or 200 ng/ml in the absence or presence of 10 ng/ml VEGF.

For luciferase assays, cells were harvested in 1× Passive Lysis Buffer (Promega E194A) for 20 min at room temperature. Firefly and Renilla luciferase activities were measured using the Dual-Luciferase Reporter Assay System (Promega E1910). Relative luciferase activity was calculated by normalizing Firefly luciferase activity to Renilla luciferase activity. All luciferase assay data were analyzed using GraphPad Prism. Data are presented as mean ± SD. All experiments were performed in triplicate.

### Western blot analysis with HEK293T cells

HEK293T cells were plated at 5×10^5^ cells/well in 6-well plates. The following day, the medium was replaced with fresh medium containing 10% serum. Three hours later, cells were transfected with the indicated plasmids using polyethyleneimine (PEI). For transfection, plasmid DNA was diluted in 200 μl serum-free medium. Cells were transfected with a total of 2.3 ug plasmid DNA/well. In a typical experiment, cells were transfected with 1 μg of receptor plasmid, 10 ng of an internal control plasmid expressing a nuclear-localized tdTomato, and either 33 ng, 100 ng, or 300 ng of Wnt1 expression plasmid, together with salmon sperm DNA to bring the total DNA to 2.3 ug. When two receptors were co-transfected (VEGFR1+VEGFR2 or TGFBR1+TGFBR2), cells were transfected with 1 μg of each receptor plasmid, 10 ng of an internal control plasmid expressing a nuclear-localized tdTomato, and either 33 ng, 100 ng, or 300 ng of Wnt1 expression plasmid, together with salmon sperm DNA to bring the total DNA to 2.3 ug. Control transfections received 2.3 μg salmon sperm DNA. DNA mixtures were combined with 9 μl PEI, incubated at room temperature for 15 min, and added to cells.

For harvesting, cells were washed three times with cold PBS and lysed in 300 μl Pierce IP Lysis Buffer (Thermo Fisher Scientific 87787) at 4°C for 30 minutes with rotating at 4°C. Lysates were centrifuged at 15,000 rpm for 10 minutes, and supernatants were mixed with loading buffer and boiled for 3 minutes.

Samples were resolved by SDS-PAGE and transferred to a Protran 0.2 um nitrocellulose membrane (Amersham 10600001) at 300 mA for 60 minutes. Membranes were incubated with primary antibodies (1:1,000 to 1:50,000) overnight at 4°C, washed, and then incubated with HRP-conjugated secondary antibodies (1:5,000). Signals were detected using SuperSignal West Pico PLUS Chemiluminescent Substrate (Thermo Fisher Scientific 34579) and a LICORbio Odyssey detector. Quantification was performed with the LICORbio Odyssey detector and software.

## Acknowledgements

Supported by the Howard Hughes Medical Institute. The authors thank David Mohr (Genetic Resources Core Facility, Johns Hopkins School of Medicine) for assistance with NextGen sequencing, Yang Li (Surrozen, Inc) for sharing the L6F4-2 amino acid sequence, and Zhongming Li for helpful comments on the manuscript.

## Author contributions

Yanshu Wang – Conceptualization, Formal analysis, Investigation, Writing - review and editing; Ningyu Zhu – Conceptualization, Formal analysis, Investigation, Writing - review and editing; Amir Rattner – Formal analysis, Writing - review and editing; Philip M. Smallwood – Formal analysis, Investigation; Jeremy Nathans – Conceptualization, Formal analysis, Supervision, Funding acquisition, Project administration, Writing - original draft, Writing - review and editing.

## Ethics

All mice were housed and handled strictly according to the approved Institutional Animal Care and Use Committee protocol of the Johns Hopkins Medical Institutions (Protocol MO25M305).

## Conflict of interest statement

J.N. is a paid consultant to EyeBio, a subsidiary of Merck, which is developing Wnt pathway activators for the treatment of retinal vascular disease.

## Supplemental Figure Legends

**Figure 4 – figure supplement 1.**
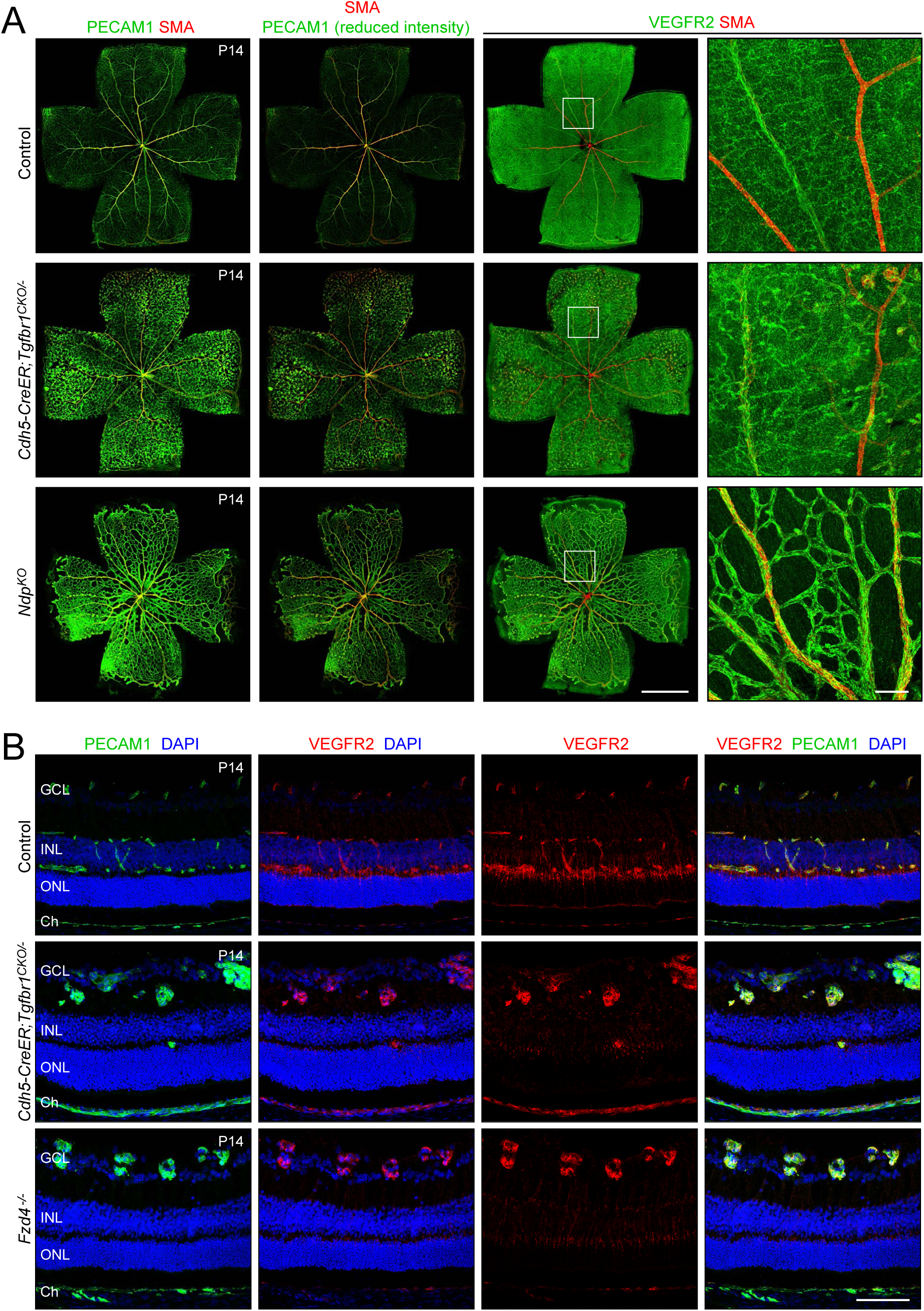
VEGFR2 levels in *Ndp^KO^* and *Fzd4^-/-^* retinas that lack endothelial beta-catenin signaling. (A) Retina flat mounts at P14 from WT control, *Cdh5CreER;Tgfbr1^CKO/-^*, and *Ndp^KO^* mice, immunostained for all blood vessels (PECAM1), for arteries (SMA), and for VEGFR2. Scale bars, 1 mm for whole retina images, and 100 um for enlarged images in the fourth column. The regions demarcated by white squares in the third column are enlarged in the fourth column. (B) Retina cross-sections at P14 from WT control, *Cdh5CreER;Tgfbr1^CKO/-^*, and *Fzd4^-/-^*mice, immunostained for all blood vessels (PECAM1) and for VEGFR2. Nuclei are visualized with DAPI. GCL, ganglion cell layer; INL, inner nuclear layer; ONL, outer nuclear layer; Ch, choroid. Scale bar, 100 um.

**Figure 5 – figure supplement 1.**
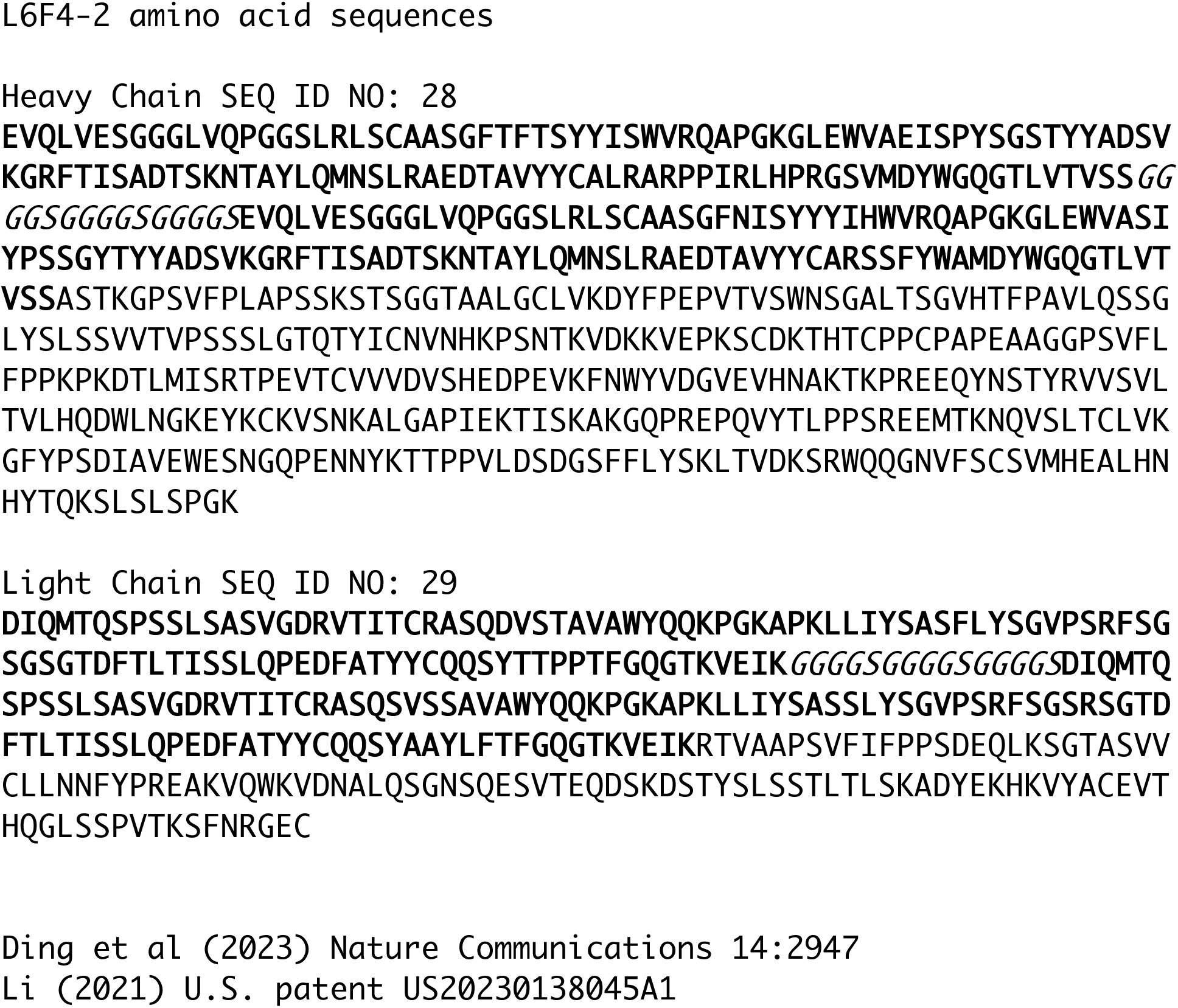
Amino acid sequence of L6F4-2. The amino acid sequences of the heavy chain (sequence ID 28) and the light chain (sequence ID 29) from Li et al (2021) U.S. patent US20230138045A1. V_H_ and V_L_ sequences are in bold. Glycine spacers are italicized.

**Figure 5 – figure supplement 2.**
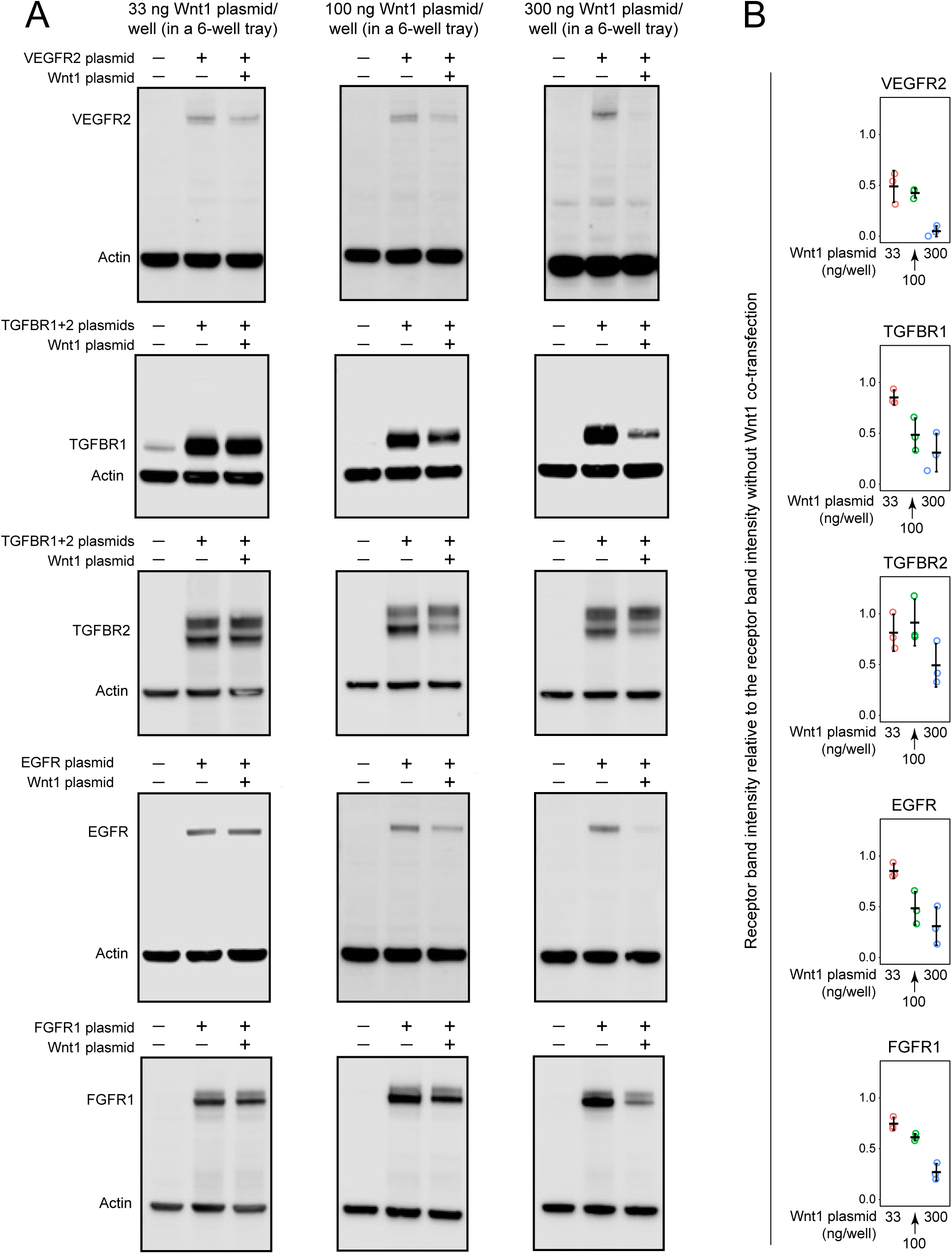
Reduced levels of VEGF, TGF-beta, EGF, and FGF receptors in HEK/293T cells with high levels of Wnt1 plasmid transfection. (A) Immunoblots of HEK/293T cells transiently co-transfected with 33 ng, 100 ng, or 300 ng of Wnt1 expression plasmid/well in 6-well plates together with 1 ug of each of the following receptor expression plasmids (from top to bottom): VEGFR2 (first row), TGFBR1+TGFBR2 (second and third rows), EGFR (fourth row), and FGFR1 (fifth row). The blots were probed with antibodies to each of the five indicated receptors. Anti-actin served as an internal control. Each blot shown here is one representative example from three independent experiments. (B) Quantification if the receptor immunoblot band intensities relative to the intensities of the samples without Wnt1 plasmid co-transfection. The data are from three independent experiments, one of which is shown in (A).

